# Reversible two-state folding of the ultrafast protein gpW under mechanical force

**DOI:** 10.1101/314583

**Authors:** Jörg Schönfelder, David De Sancho, Ronen Berkovich, Robert B. Best, Victor Muñoz, Raul Perez-Jimenez

## Abstract

Ultrafast folding proteins have limited cooperativity and thus are excellent models to resolve, *via* single-molecule experiments, the fleeting molecular events that proteins undergo during folding. Here we report single-molecule atomic force microscopy (AFM) experiments on gpW, a protein that, in bulk, folds in a few microseconds over a marginal folding barrier (~1 *k*_B_*T*). Applying pulling forces of only 5 pN we maintain gpW in quasi-equilibrium near its mechanical unfolding midpoint, and detect how it interconverts stochastically between the folded and an extended state. This binary pattern indicates that, under an external force, gpW (un)folds over a significant free energy barrier. Using molecular simulations and a theoretical model we rationalize how force induces such barrier in an otherwise downhill free energy surface. Force-induced folding barriers are likely a general occurrence for ultrafast folding biomolecules studied with single molecule force spectroscopy.

---

Deciphering the mechanisms by which proteins fold has long been one of the central problems in molecular biophysics^1,2^. This quest has proved challenging because most single domain proteins fold slowly via a two-state (i.e. all or none) process^3^, and atomistic simulations could only access very short timescales^4^. In this context, downhill folding attracted particular attention with the promise of unveiling details of folding energy landscapes that are hidden in two state folding^5^. Downhill folding proteins do not cross significant free energy barriers and thus exhibit limited cooperativity^6^ and are amongst the fastest to fold and unfold^7^. Their μs folding times have been instrumental in bridging the time scale gap between experiment and atomistic molecular dynamics (MD) simulations^7-11^. The minimal cooperativity of downhill folding has led to methods that distil mechanistic information from conventional ensemble experiments, such as monitoring how thermal denaturation depends on the structural probe^12^, analyzing heat capacity thermograms in terms of low-dimensional free energy surfaces^13^, or estimating free energy barriers to folding from the curvature of the Eyring plot^14^.

Whereas many fast folding proteins share common structural features like their small size (typically, less than 45 residues) or primarily helical secondary structure (with the exception of the very small WW domains), the protein gpW is an outlier to these general trends^15^. gpW has 65 residues and a native α+β structure that consists of two antiparallel α–helices and a single antiparallel two-stranded β-sheet, but it folds and unfolds in only ~4 μs at the denaturation midpoint and exhibits the characteristic features of downhill folding^15^, including minimally cooperative (un)folding that results in many different patterns at the atomic level when investigated by Nuclear Magnetic Resonance (NMR)^16^. Both atomistic MD simulations^16^ and simple statistical mechanics models^15,17^ agree in classifying gpW as a downhill folder. These properties make this protein an attractive candidate for smFS studies, which to our knowledge have been previously conducted for just two other ultrafast folders (villin^18^ and α3D^19^).

Here we employ an Atomic Force Microscope (AFM) that allows us to make stable measurements at low forces (between 3 and 10 pN) in the constant force mode^20,21^. Surprisingly, gpW behaves in these experiments as a reversible two-state folder, with clearly distinguishable hopping events between the native and an extended state. Detailed analysis of the experiments, coarse grained molecular dynamics simulations and a theoretical model, indicate that the pulling force induces a free energy barrier to the (un)folding of gpW, thus confirming experimentally the scenario of force-induced refolding barriers observed in molecular simulations of RNA^22^ and predicted for protein unfolding^23^.

## Results

### Force ramp experiments reveal the mechanical unfolding midpoint

To measure the mechanical unfolding of gpW using the AFM we designed and expressed a polyprotein construct where gpW was sandwiched between three titin I91 domains of the human cardiac protein on each side (Fig. 1a). In this construct, the titin I91 domains serve as molecular fingerprint for AFM trace selection due to their well characterized unfolding force and contour length^24^. We performed force-ramp measurements using this construct to determine the mechanical stability of gpW. In these experiments, we ramped the force very slowly (1 pN s^−1^), starting from pushing the AFM tip against the surface with a 10 pN force (*F*<0), gradually moving to the pulling regime, and ending at a final pulling force of 150 pN (*F*>0) so that we could unfold the six I91 domains. The recorded traces showed a first extension event of ~10-11 nm that takes place at times varying between 10 and 25 seconds (i.e. 0 to 15 pN). These are followed much later by six, sharp ~24.5 nm unfolding events, the extension expected for each of the I91 titin domains in the polyprotein (Fig. 1b).

**Fig. 1.**
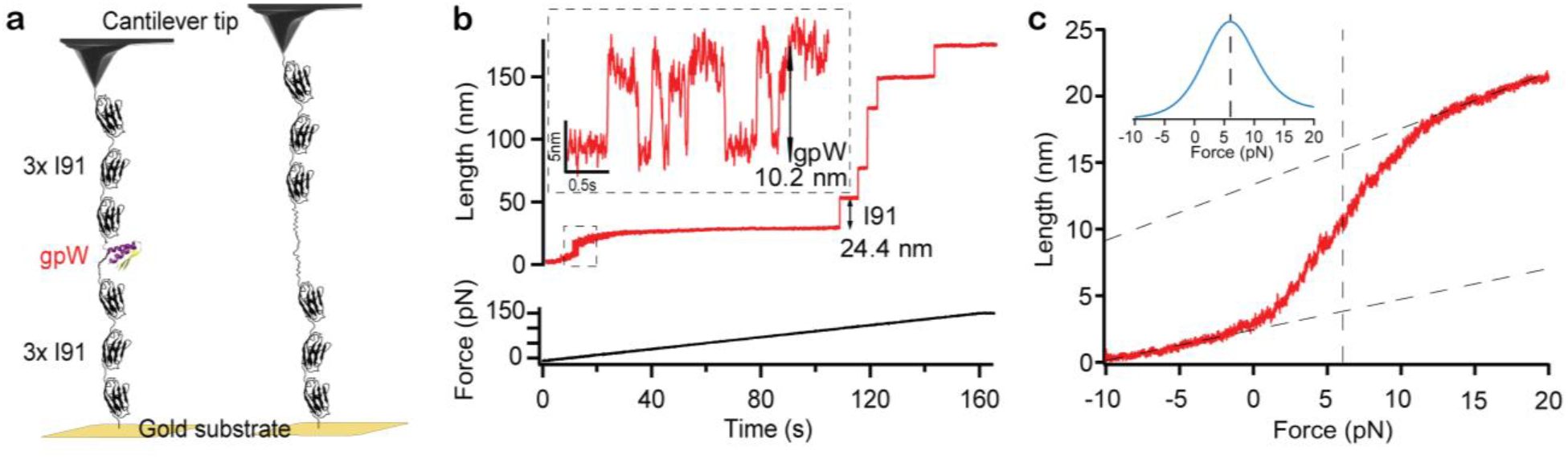
GpW unfolds mechanically at small forces. (a) Schematic of the polyprotein construct (I91)3-gpW-(I91)_3_. The sample is adsorbed to the gold substrate on one end and to the AFM cantilever on the other. Upon application of a pulling force below 15 pN gpW interconverts between its extended unfolded and native state. (b) AFM force ramp trace of gpW at a velocity of 1pN s^−1^ showing the titin I91 fingerprint at the end of the trace. Insets show the hopping pattern of gpW occurring at low forces. (c) Average length vs force plot at a force ramp of 1 pN s^−1^ (29 force ramp traces) showing the mid unfolding force of gpW at around 6 pN. Inset shows the derivative of the curve. Propagating baselines lead to an estimated extension for gpW of ~10.5 nm.

Based on the change of extension associated to the isolated first event (~10 nm), we can tentatively assign it to the unfolding of gpW. Unfolding at low forces is expected for a largely α– helical protein that shows marginal stability in ensemble chemical denaturation experiments^15^. Moreover, a ~10 nm extension is commensurate with the expectation for a worm-like chain of 65 residues at these very low forces (see below). Interestingly, this extension event is not a single step. In fact, zooming into the low force region in our traces reveals hopping patterns that are consistent with stochastic series of extension and retraction steps (~10-11 nm). Such hopping continues until the cantilever settles at the 10-11 nm extension when the force raises above 15 pN. These observations can be interpreted in terms of multiple folding-unfolding interconversions of gpW taking place at the very low ramping rate of these experiments. The interconversions keep on occurring until the force reaches values high enough to maintain it mechanically unfolded. Past this point the traces remain flat for very long times (>80 seconds) highlighting the stability of our AFM.

To determine the midpoint unfolding force of gpW, we averaged 29 force ramp curves similar to that shown in Fig. 1b and which contained the mechanical fingerprint of at least 4 I91 domains. Averaging was performed in the force regime between 10 pN pushing (*F*<0) and 20 pN pulling (*F*>0). The resulting length *vs* force curve represents the cumulative distribution of unfolding forces for gpW (Fig. 1c). The derivative of this curve has its maximum at ~6 pN (inset to Fig. 1c), indicating the mid-unfolding force of gpW. Propagating the low and high force baselines to the center of the sigmoid results on an estimated extension of 10.5 nm for gpW at 6 pN that matches perfectly the prediction from the worm-like chain (WLC) model25 for a polymer with the properties of gpW: persistence length ρ~ 0.8 nm^26^ and contour length *Lc* ~ 23.4 nm (our used gpW protein has 65 residues including tails and the crystallographic contour length for an amino acid is 0.36 nm) and a N-C termini distance of around 1nm (derived from the crystal structure of gpW pdb file 2L6Q ^27^).

### Force control of the folding free energy landscape of gpW

The spring constant of the Biolever cantilevers that we use is around 5 pN nm^−1^, which precludes more accurate estimates of the midpoint force using the AFM in the force-clamp mode. However, we could verify that the apparatus can shift the mechanical unfolding equilibrium of individual gpW molecules towards the folded or the unfolded state using experiments that combine force-ramp and force-clamp AFM measurements (Fig. 2). Particularly, we switched between three applied constant forces in the low force regime (0 – 15 pN) using slow force ramp segments (1 pN s^−1^), followed by a final increasing force ramp (from 1 to 20-100 pN s^−1^) to quickly unfold the 6 flanking titin I91 domains. We used two alternative force routines. In one of these, the force is switched from 5 pN (roughly the midpoint) to 10 pN (mostly unfolded), and then back to 5 pN (Fig. 2a). The second routine changes the constant force in steps of 3, 5 and 8 pN (Fig. 2b). In both types of experiments we observe shifts in the equilibrium populations between the native state and a mechanically unfolded state that reflect the tilting of the gpW folding landscape through the application of a pulling force. However, the quantitative interpretation of this data needs to consider how the spring constant of the cantilever affects the experimental resolution in force. As it can be seen in Fig. 2a,b, the force fluctuates with a standard deviation (SD) of ±2 pN. Using the WLC model with the parameters discussed above, a 2 pN change in force corresponds to a change in length of 2-3 nm in the low force regime (i.e. around 5pN). In Fig 2a, for example, the difference in extension between the 10 and 5pN segments is around 3nm, which is consistent with the WLC model. In contrast, in Fig 2b. the unfolding length of gpW during the 3pN and 5pN segments is very similar because their difference is within the force resolution limit of the cantilever, which cannot measure force differences < 4 pN (2x SD). Here we need to mention that we are operating in a force regime that is much lower than in conventional AFM experiments. This limitation combined with the fact that we maintain the force constant for more than 1 minute, make these experiments extremely challenging and unique. Nevertheless, from the experiments shown in Fig.2 we can phenomenologically conclude that the small applied forces modulate the unfolding behavior of the protein.

**Fig. 2.**
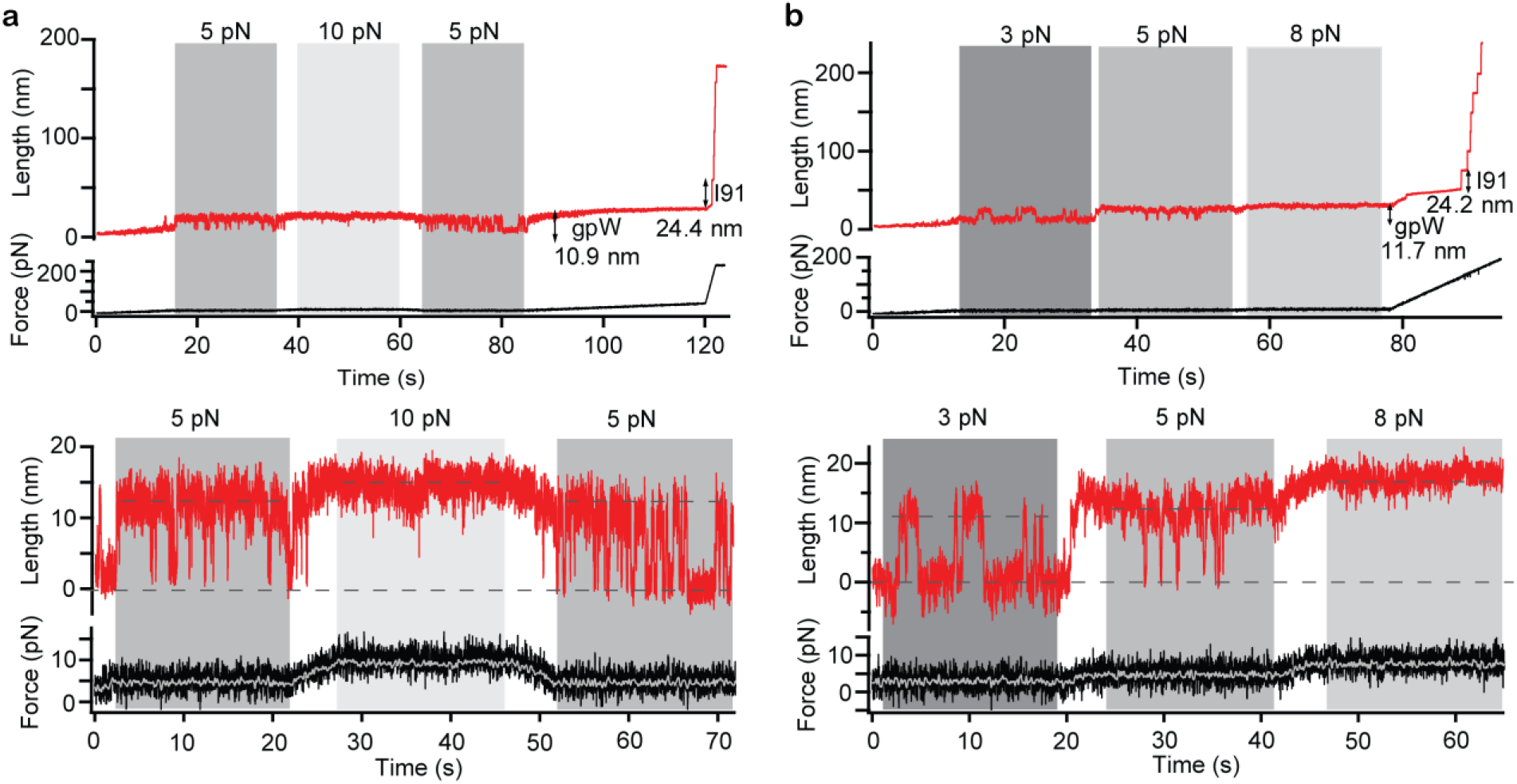
Mechanical force modulates the (un)folding equilibrium of gpW. Length *vs*. time (top) and force vs. time (bottom) from experiments were conducted using a force sequence of 5 pN-10 pN-5 pN (a) and 3 pN-5 pN-8 pN (b). Dashed black lines indicate the extension at each applied force. The force vs. time trace indicates the force resolution of the used Biolever AFM cantilever, showing both the digitally filtered force (grey) and unfiltered signal (black).

### Two-state hopping behavior in constant force experiments

We next performed force-clamp AFM measurements at 5 pN to analyze in more depth the folding-unfolding of gpW around its mechanical denaturation midpoint. As before, we designed an experiment that starts with a force-ramp segment at 1 pN s^−1^ to reach 5 pN, continues with 20-30 seconds in which the force is kept constant at 5 pN, and ends with a force ramp segment that hikes the force at an increasing rate (from 1 to 20 pN s^−1^). We collected 13 such experimental traces that also showed the mechanical fingerprint for at least 4 I91 domains. These traces reveal the distinct patterns of alternating ~10 nm extensions and retractions that indicate the reversible mechanical unfolding-refolding of gpW. We show one such trace in Fig. 3a. Five more traces are superimposed in Fig. S1 to illustrate the reproducibility of these experiments.

**Fig. 3.**
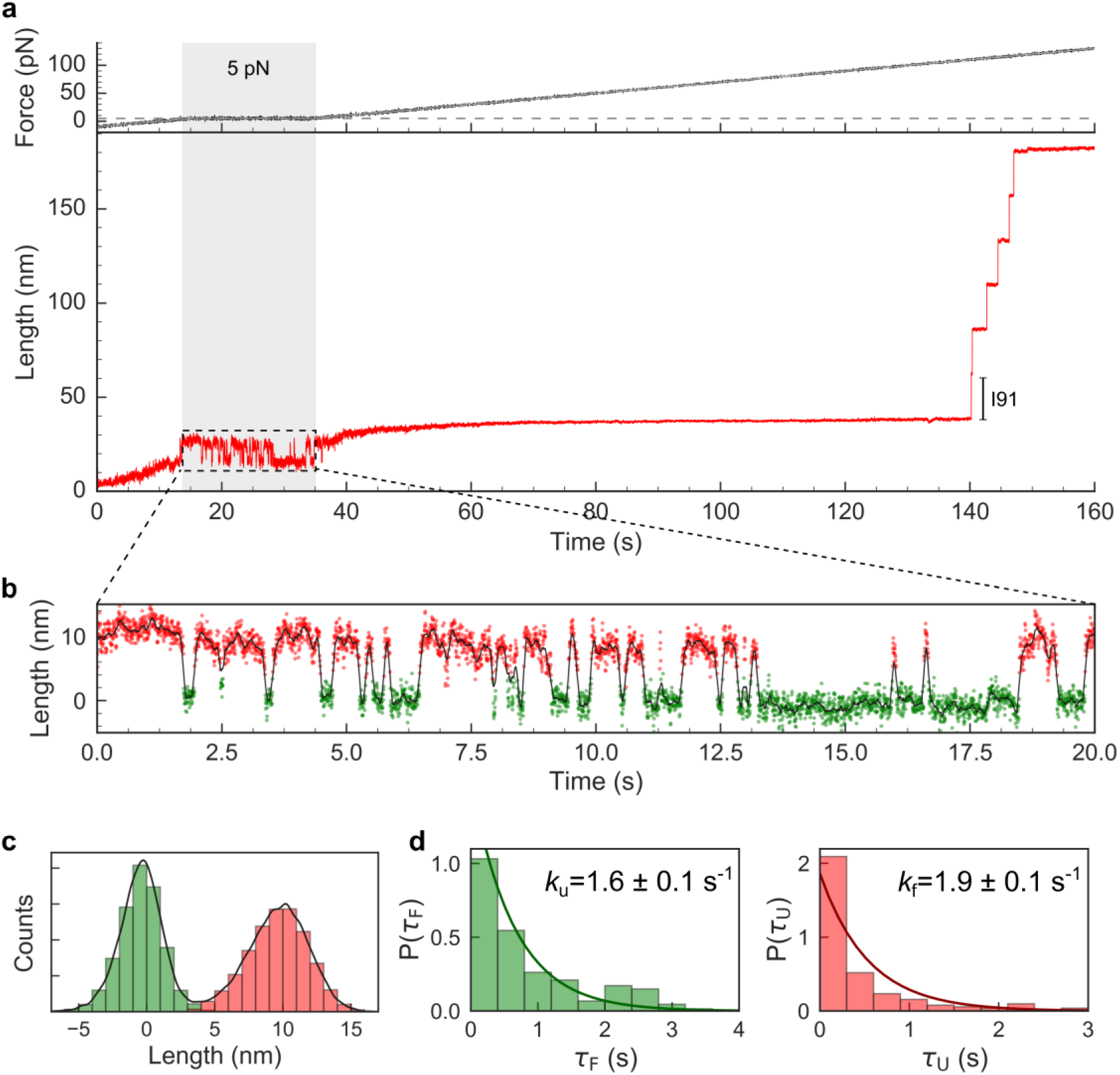
Force clamp experiments on gpW and lifetime analysis. (a) A complete force (top) and length (bottom) *vs*. time trace from our AFM force-clamp measurements. gpW unfolds and refolds in the 20 s segment held at a constant 5 pN force (marked by a grey swath) and the six I91 domains unfold at the end of the trace. (b) Detail of the 5pN segment of the length *vs* time trace revealing the hopping pattern of gpW with extensions retractions of ~10.0 nm. The folded state is colored in green and the unfolded state in red, based on the assignment from a hidden Markov model. (c) Histograms of lengths for the folded and unfolded states at 5 pN and corresponding probability distribution (black line). (d) Distribution of lifetimes in the folded (green) and unfolded (red) states. In both cases the lifetime histograms were fitted to an exponential distribution (lines).

Zooming into the 5 pN constant force segment of these curves (Fig. 3b) reveals the stochastic nature of the folding-unfolding events. To make our observations quantitative, we analyzed all the traces using the PyEMMA software package to generate a hidden Markov model (HMM) from the time series data of measured extensions (see Methods and Fig. S2). The HMM analysis supports the definition of two unique states given the time-scale separation between the slowest mode (folding / unfolding) and fast dynamical processes (see Figure S2c). Using the trajectories assigned with the HMM we can produce the histogram of molecular extensions, which shows a bimodal distribution with one peak at ~0 ± 2 nm (native) and a second peak at ~10 ±3 nm (unfolded) (Fig. 3c). The populations for the unfolded and native states obtained from the length histogram at 5 pN force are nearly equal, confirming that 5 pN is in fact very close to the mechanical denaturation midpoint for gpW (Fig. 3c), as we estimated from the force-ramp experiments (Fig. 1c). From a simple Boltzmann inversion of the distribution of molecular extensions we can obtain a free energy surface for the mechanical unfolding of gpW as a function of extension. This surface presents a folding free energy barrier of ~2.5 *k*_B_*T* between the native and extended unfolded ensembles that corresponds to a lower bound for the actual barrier.

We determine the lifetimes of gpW in the native folded and in the mechanically unfolded states from the distribution of dwell times in the HMM states. These distributions are well fitted to single exponential functions with rates constants of 1.9 ± 0.1 s^−1^ for refolding (red in Fig. 3d) and 1.6 ± 0.1 s^−1^ for unfolding (green in Fig. 3d). These timescales are much slower than the response time of our instrument, which in these experiments is equivalent to the sampling rate since our piezo electric actuator completes the force compensation through the feedback loop in times <1 ms (Fig. S3). Additionally, the response time of the cantilevers we use is in the range of 0.05-1 ms^28^. Interestingly, the dynamics of the folding-unfolding transitions under mechanical force is also very slow relative to the folding and unfolding rates of ~60,000 s^−1^ that have been measured for untethered gpW at the midpoint temperature (343 K). In fact, the slowdown is so drastic that, if it were caused solely by an induced folding barrier, it would have to be exceedingly large (~10 *<u>k</u>*_B_*T*). This observation suggests that much of the slow-down may be caused by dynamic effects of the measuring device^29,30^. An analysis of such instrument-based effects is beyond the scope of this work, and will discussed in a follow up publication. Nevertheless, we can confirm the shift of gpW from downhill folding in the absence of force to a two-state folding regime under mechanical force without invoking the slowness of the observed rates. The two-state nature of the process is indeed apparent in several complementary observations: 1) the single exponential distribution of lifetimes; 2) the distribution of molecular extensions; 3) the binary switching patterns observed in the experimental traces; 4) and the results from the HMM.

### Coarse grained molecular simulations reproduce the force-induced barrier

To rationalize our experimental results, we performed molecular simulations with a coarse-grained, structure-based model^31^ that has been used before to describe mechanical unfolding experiments^32,33^. We parameterized the model with the recently determined 3D structure of gpW^27^ and ran simulations pulling from the ends of the protein at different forces. When we project the simulation data on the pulling coordinate *x* (the end-to-end distance), the resulting traces show two-state hopping patterns that bear close resemblance to experiment (Fig. 4a). Simulations in the presence of force lead to a highly extended unfolded state (see Fig. 4b). The difference in extension between the folded and unfolded states in the simulation (~6 nm) is not in quantitative agreement with the experimental value. This discrepancy is likely due to an unrealistically high intrinsic helical propensity in the simulation model, which maintains the two helices fully formed under 6 pN force. However, the general (un)folding behavior is correct in qualitative terms. In Fig. 4c we show the potential of mean force (PMF) for the projection on the molecular extension (*x*) in the absence of force, and at the simulation mechanical midpoint (6 pN). Comparison of the two PMFs reveals a downhill free energy landscape in the absence of force, and the emergence of a force-induced free energy barrier that separates the folded and unfolded states at 6 pN. Two-dimensional PMFs obtained by umbrella sampling at different forces (Fig. 4d) show that the pulling coordinate (*x*) and the folding coordinate (the fraction of native contacts, *Q*) are correlated even at the lowest forces. Therefore, contrary to previous findings for the two-state folder GB1^33^, the downhill character of gpW in the absence of force is not an artifact arising from the use of the molecular extension as order parameter for folding in the no-force regime. In the simulations, the barrier induced by force is 5.3 kJ mol^−1^ (or ~1.7 *k*_B_*T*) for unfolding and 4.8 kJ mol^−1^ (or ~1.6 *k*B*T*) for refolding. These relatively lower free energy barriers observed in the simulations compared to experiments are expected considering that coarse-grained structure-based models generally underestimate the cooperativity of protein folding^34^. We obtained similar results in simulations with a phenomenological one-dimensional model specifically developed for mechanical unfolding^35^ (Fig. S4). The latter model also permitted to simulate the complex force routines of Fig. 1, which are again in semi-quantitative agreement with experiment.

**Fig. 4.**
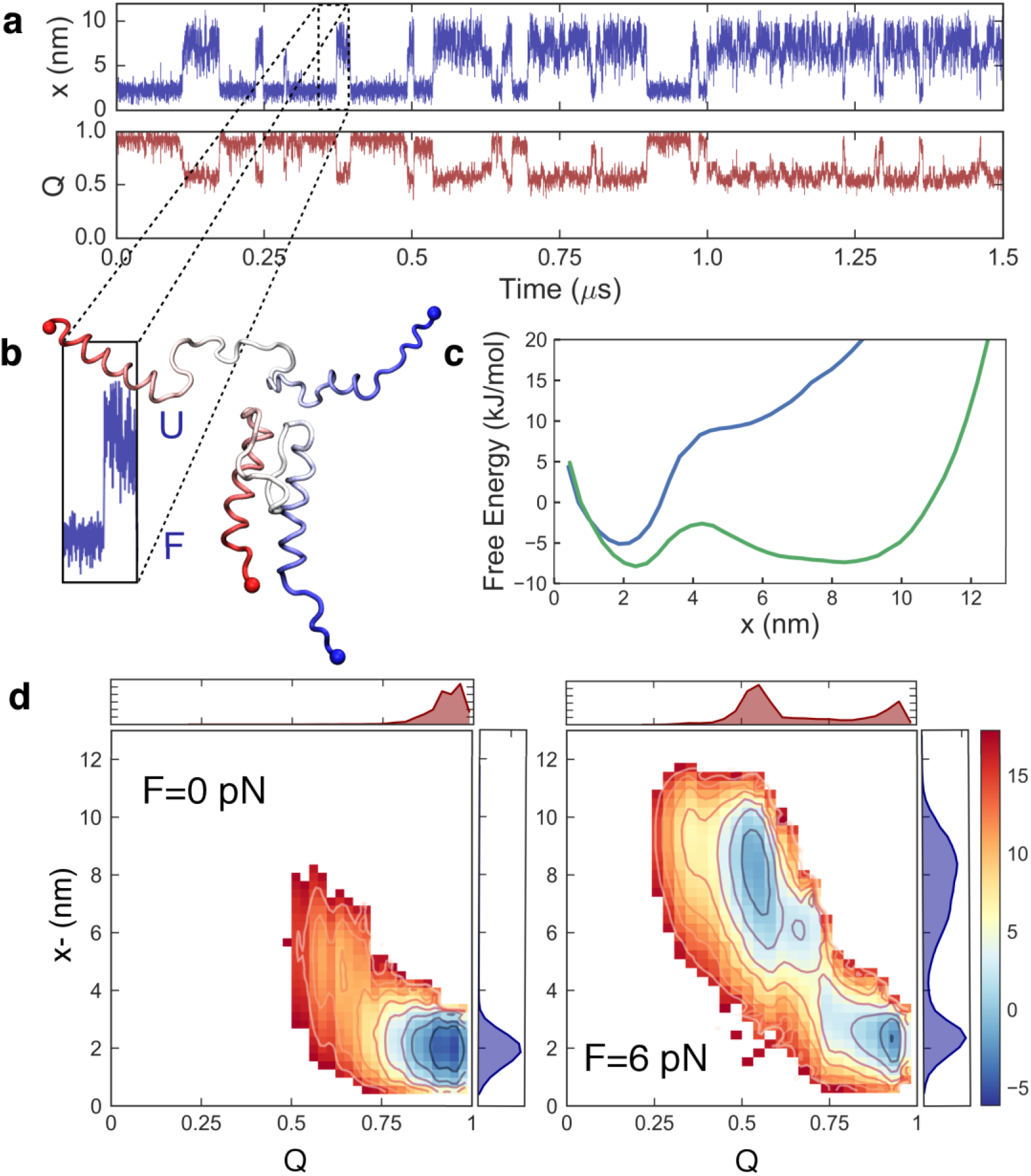
Coarse grained structure based molecular simulations of gpW. (a) Time series data from a constant force simulation at 6 pN projected on the end to end distance, *x*, and the fraction of native contacts, *Q*. (b) Blown out segment of the time series on *x* and representative snapshots corresponding to the folded and unfolded states from the forced unfolding simulations. (c) Free energy profiles for the projection on the end to end distance (x (nm)) at 0 and 6 pN. (d) Two dimensional potentials of mean force on the fraction of native contacts (*Q*) and the end to end distance (*x*) at 0 (left) and 6 pN (right). Top and side plots indicate equilibrium populations for the one dimensional projections on *Q* (red) and *x* (blue). Free energies are expressed in kJ mol^-1^.

The reasonable compliance between experimental results and molecular simulations of single-molecule pulling^36^ is noteworthy, especially considering the simplified nature of the simulations. Such general agreement gives us license to infer certain mechanistic details from the simulations. The mechanism that emerges is one in which force gradually peels off the two α-helices, starting from the termini. The barrier top is reached at the point in which only the last segment of H1 and the first of H2, together with the loop hinges connecting them to the hairpin, remain in contact (Fig S5). When the tertiary contacts between these two structural segments break, the protein unravels; maintaining the helices formed in simulations, and probably with the helices unfolding more prominently in experiments. The mechanical unfolding of the α+β protein gpW is thus very different from that of the all-β CspB, in which the 6 β-strands unravel one by one stochastically21. Similar mechanistic differences have been found for these two proteins on a recent computational analysis of folding pathways^37^. It is interesting to note that the mechanical unfolding transition state of gpW that we find here and the thermal unfolding transition state inferred from the folding interaction networks obtained by NMR experiments and MD simulations^16^ appear to be quite similar. The commonality between the anisotropic unfolding by pulling and thermal denaturation suggests that what makes proteins (un)fold ultrafast, and particularly the relatively large α+ β protein gpW^15^, is their folding occurring via a diffuse mechanism characterized by broad ensembles of parallel microscopic pathways.

## Discussion

Here we have determined the mechanical stability of the ultrafast folder gpW using an AFM that operates at very small loads (3-6 pN). When subjected to a constant pulling force that tilts its folding free energy landscape to the mechanical denaturation midpoint (~5 pN), gpW undergoes remarkably slow interconversions (~4-5 per second) between its compact native state and a stretched unfolded ensemble. The stochastic switching patterns of gpW are perfectly amenable to a two-state description, in contrast with the continuum of states that one might expect for a barrierless (downhill) protein.

Over the last years, several studies have reported the detection of conformational transitions of mechanically controlled proteins^38-41^. Similar conformational transitions have been reported for another ultrafast folder, the villin headpiece subdomain^18^. Constant position, equilibrium measurements with an AFM equipped with improved cantilevers have recently unveiled the complex unfolding pathways of bacteriorhodopsins^42^. However, our results on gpW represent the first example of measurements at quasi-equilibrium of individual protein molecules interconverting between their native and mechanically unfolded states using force-clamp AFM. This was possible because our AFM allows for stable measurements performed at relatively small constant forces, over extended periods of time, with a relatively fast time response, and using simple protein constructs.

Our results highlight the different nature of thermal and mechanical denaturation^32^ and demonstrate that the scenario of force-induced barriers proposed by Fernandez and coworkers does indeed apply to downhill proteins, which lack a free energy barrier in the absence of force^23^. The key to these results resides in the polarizing nature of the applied force, which selectively stabilizes highly extended unfolded conformations that would be rarely populated in the absence of force. At low force the unfolded ensemble is kept relatively compact by conformational entropy, but as force increases, the ensemble becomes mechanically compliant by selecting extended conformations over other, more compact, unfolded conformations. As a consequence, the unfolded minimum in the folding free energy landscape moves far apart (in terms of end-to-end distance) from the native state so that a force-induced free energy barrier emerges between the two minima, even if the protein (un)folds downhill in the absence of force.

## Methods

### Cloning and Protein Expression

The chimeric polyprotein construct (I91)_3_-gpW-(I91)_3_ was designed containing the gpW protein (Top Gene Technologies, Canada) flanked by three Titin-I91 domains on each side using standard DNA manipulation protocols to build the construct inside the pRSET A vector. Each DNA manipulation step needed to add a protein domain consecutively into the plasmid vector was confirmed by DNA sequencing (Parque Científico, Madrid). C41 strand competent cells *E. coli* were used for protein expression as they are specialized in expressing toxic proteins (Novagen). A gentle cell lysis protocol was used to avoid damage to the expressed polyproteins^43^. The sample was then purified by HPLC (Agilent, Santa Clara, CA) in two steps: first using a nickel-affinity HisTrap column (Ge Healthcare) and second using a size exclusion Superdex 200 column (GE Healthcare). Finally, the buffer was changed to the measuring buffer 1x PBS pH 7.4 using ultrafiltration Amicon 3k filters (Milipore). The final protein concentration was estimated to be around 1 mg ml^−1^ using a Nanodrop (Thermo Scientific). Then the samples were snap frozen in liquid nitrogen and stored at −80°C.

### Single Molecule Force Spectroscopy

All single-molecule force spectroscopy constant force and force ramp experiments were performed on a force clamp AFM (Luigs Neumann) ^44^. Biolever cantilevers from Olympus/Bruker were used with a spring constant 3 - 8 pN nm^−1^ for all constant force and force ramp measurements. The spring constant was measured before each experiment using the equipartition theorem within a software built-in procedure. Data was recorded between 0.5 to 4 kHz for the constant force and force ramp measurements. During combination of force-ramp and constant force experiments, the force was ramped at rate of 1pN s^−1^ until reaching the 5 pN constant force (starting from 10 pN pushing *F*<0). Then the protein was held for 20-30 s at the constant force before the ramp at 1pN s^−1^ was continued until the six Titin-I91 domains were unfold. For the force-clamp experiments with multiple applied constant forces we ramped the polyprotein construct until the desired constant force at a rate of 1 pN s^−1^. After holding the protein under the constant force for 5-20 s we switched to another force using again a force ramp of 1 pN s^−1^. After applying all desired constant force segments we applied a force ramp of 1 pN s^−1^ until the six Titin-I91 domains were unfolded. To reduce total experimental acquisition time of force-clamp experiments, we ramped at 1 pN s^−1^ up to reaching 60 pN, and then increased the force rate to 20-100 pN s^−1^ to quickly reach the high forces required to unfold I91. In the force ramp experiments the force was ramped at a rate of 1pN s^−1^ throughout the whole experiment up to a value of 150pN (starting from 10 pN pushing *F*<0) at which the six Titin-I91 were unfolded.

### Experimental Conditions

All AFM experiments were carried out at room-temperature (~24 °C) in 1x PBS buffer at pH 7.4. Typically, 40 μl of the protein sample (~μM concentration) was left around 20 minutes for adsorption on a fresh gold coated surface (Arrandee). After the adsorption time the sample was then rinsed of the gold surface by 1x PBS buffer to remove unbounded protein sample just before starting the measurements.

### Data Analysis with a Hidden Markov model

Data from AFM experiments were first screened and analyzed in Igor Pro (Wavemetrics) using the built-in data analysis procedure file and then with Python in-house scripts. First, we shifted all the trajectories so that the native state is centered at *x* = 0. Then we assigned the trajectories to two states using a Hidden Markov model (HMM) built from the experimental traces with the PyEMMA package^45^. The procedure involves the generation of a fine-grained Bayesian Markov state model, for which we binned the extension data into 50 microstates. This model was validated using both the convergence of the implied time-scales for the slowest relaxation time of the system and a Chapman-Kolmogorov test. From this fine-grained model we constructed a hidden Markov model. The separation of time-scales of ~1 order of magnitude between the slowest mode and the faster modes in the fine-grained model affords a separation into two unique states. Given that we had collected data at two different sampling frequencies, we carried out the analysis of the two datasets separately. Finally, from the assigned trajectories we computed the lifetimes for the folded and unfolded states, and fitted them to a single exponential distribution.

### Coarse grained molecular dynamics simulations

#### Structure-based model

We run simulations of gpW using the Karanicolas and Brooks structure-based (i.e. Gō) model^31^. In the simulations, the protein geometry is reduced to the C^α^ trace and the solvent degrees of freedom are not considered, resulting in a great computational efficiency. The potential energy is calculated as

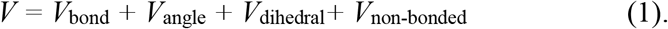

In Equation 1, *V*_bond_ and *V*_angle_ are native-centric harmonic terms for bonds and angles, respectively, while *V*_dihedral_ is based on statistical preferences for torsion angles in the PDB^31^. The non-bonded contribution is favourable for pairs of residues that are in contact in the reference (i.e. experimental) structure. Two residues *i* and *j* are defined to be in contact when any pair of atoms is closer than 5 Angstroms, and their contribution to the energy is

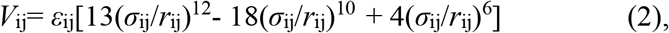

where *σ*_ij_ is the distance between the C^α^’s of residues *i* and *j* in the reference structure, *r*_ij_ is the same distance but in the instantaneous configuration and *ε*_ij_ is the strength of the pairwise interaction^31^. We used the recently determined experimental structure of gpW (PDB id: 2l6q27) as reference. In order to recover a melting temperature comparable to that from the experiments (340 K^15^) we scaled up the native contacts by 50%.

#### Molecular simulations

Simulations were run using a modified version of Gromacs 4.0.5^46^. We propagated the dynamics at a constant temperature of 300 K using a Langevin integrator with a time-step of 10 fs, and a friction constant of 0.2 ps^−1^. Pulling experiments were simulated using the pull-code from Gromacs by defining pulling groups in the protein ends and pulling in a single dimension at constant force values between 0 and 10 pN. Additionally, for the calculation of potentials of mean force, we run umbrella sampling simulations at different force values by imposing umbrella potentials at equally spaced intervals on the fraction of native contacts (*Q*). The data from the simulations were projected on different progress variables, including the end to end extension relevant to the single molecule pulling experiments, and the fraction of native contacts (*Q*), defined as before^47^. The results from the umbrella sampling runs at different values of the umbrella coordinate were combined using the weighted histogram analysis method^48^. From the constant force runs we estimated kinetics by simply imposing a cutoff value for each of the states.

### Brownian Dynamics on an empirical model

We performed BD simulations following the procedure described before using a naïve representation of the PMF profile^49^. This potential is written down as the sum of attractive excluded volume interactions given by a Morse potential, an entropic term described by the worm-like chain model of polymer elasticity, and a force-dependent term that reflects the effect of the applied mechanical load on the system. Over this potential, we collected position time series by numerically solving the over-damped Langevin equation, which is a momentum balance given by

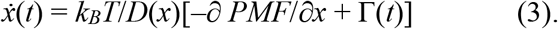

Here *D*(*x*) is the diffusion coefficient taken as 100 nm^2^/s in the folded state and 4000 nm^2^/s when the protein unfolds (crosses the barrier), and Γ(*t*) is a thermally fluctuating random force with zero mean and amplitude given by the fluctuation-dissipation theorem, (2*Ddt*)^1/2^. The PMF was characterized with an energy barrier of 3 *k_B_T* located at 2.7 nm from the native state basin. For the elastic term, we used a persistence length of 0.8 nm and gpW contour length of 23.4 nm.

## Acknowledgements

DDS and RBB wish to acknowledge Attila Szabo (NIH-NIDDK) for many helpful discussions. This research has been funded in part by the European Research Council (grant ERC-2012-ADG-323059 to VM). DDS acknowledges a grant (CTQ2015-65320-R) and Ramón y Cajal contract (RYC-2016-19590) from the Spanish Ministry of Economy and Competitiveness. RB acknowledges the support of the I-CORE Program of the Planning and Budgeting Committee and The Israel Science Foundation (Grant No. 152/11). RBB was supported by the intramural research program of the National Institute of Diabetes and Digestive and Kidney Diseases of the National Institutes of Health. VM acknowledges additional support from the W.M. Keck Foundation, the CREST Center for Cellular and Biomolecular Machines (grant NSF-CREST-1547848) and the NSF (grant MCB-1616759). RPJ acknowledges support from the Spanish Ministry of Economy and Competitiveness (grant BIO2016-77390-R) and European Commission CIG Marie Curie Reintegration program FP7-PEOPLE-2014.

## Author contributions

JS and DDS contributed equally to this work. JS performed all experiments and analyzed the data. DDS performed the coarse-grained simulations and analyzed both the experimental and simulation data. RBB contributed materials and analyzed the data. RB ran the Langevin simulations. VM and RPJ conceived the project and supervised the work. All the authors contributed to interpreting the results and writing the manuscript.

